# FungiGutDB: Curated database for taxonomic assignment of the gut mycobiome by whole genome sequencing

**DOI:** 10.64898/2025.12.03.691829

**Authors:** Diego Coleto-Checa, Blanca Lacruz-Pleguezuelos, Alba Perez-Cuervo, Nicolas Cárdenas-Roig, Lucía Carrasco-Guijarro, Adrián Martín-Segura, Enrique Carrillo de Santa Pau, Laura Judith Marcos-Zambrano

**Affiliations:** Computational Biology Group. IMDEA Nutrition, CEI UAM+CSIC. Carretera de Cantoblanco, 8. 28049 Madrid (Spain); UAM Doctoral School, Universidad Autónoma de Madrid, Spain

**Author notes:** Address correspondence to Laura Judith Marcos-Zambrano, Enrique Carrillo de Santa Pau.

**Keywords:** gut mycobiome, database, microbiome, fungi, health, nutrition

## Abstract

Fungi represent less than 1% of the gut microbiota; However, their importance in host homeostasis and disease is increasingly recognized. Accurate characterization of the gut mycobiome from metagenomic data remains a significant challenge due to the low abundance of fungal DNA, the performance of bacteria-oriented classifiers, and the limited availability of curated fungal reference databases. To overcome these issues, we developed FungiGutDB v1.0, a curated database containing 304 taxa previously identified in culture-dependent human studies, and we integrated the database in a reproducible workflow to ease its application (FungiGut).

Benchmarking analyses demonstrated that FungiGut achieved a substantially lower false positive rate in mock communities compared to standard non-gut-specific fungi databases. When applied to real metagenomic datasets, FungiGut successfully characterized the gut mycobiome, identifying Saccharomyces cerevisiae as the predominant species in healthy individuals, along with common dietary fungi found in fermented dairy products (*Penicillium camemberti, Debaryomyces hansenii, Kluyveromyces lactis, Pichia kudriavzevii*). In contrast, samples from patients with non-responsive celiac disease showed a higher relative abundance of opportunistic pathogens and a lower number of diet-associated taxa, suggesting a trend toward a dysbiotic mycobiome profile.

By limiting classification to fungal species previously isolated from the human gut, FungiGut minimizes misclassifications derived from environmental or plant-associated taxa, which often lead to mistaken interpretation of the results. Overall, FungiGut offers a biologically consistent and reproducible approach to gut mycobiome profiling, improving taxonomic accuracy and strengthening confidence in the interpretation of fungal metagenomic data in human microbiome research.

## Introduction

The importance of the gut microbiome in human health is well established, with key roles in metabolism and host physiology. However, most research has focused primarily on bacteria, while the contribution of fungi has received comparatively less attention. Although the fungal fraction of the gut microbiome represents less than 1% of the total microbial community, it plays an important role in maintaining host homeostasis and influencing both physiological and pathophysiological processes [1].

The majority of studies characterizing the gut mycobiome rely on Internal Transcriber Spacer (ITS) amplicon sequencing, a cost-efficient method that nonetheless has limited taxonomic resolution [2]. ITS regions can be highly similar among closely related species, leading to misclassification or restriction to genus-level assignments. Moreover, performance depends on multiple factors, including the choice of ITS subregion (ITS1, ITS2, or full-length), the completeness and curation of reference databases, the bioinformatic pipeline, and the fungal groups under study [3], [4]. In contrast, shotgun metagenomic sequencing offers improved taxonomic resolution and enables the simultaneous detection of fungal, viral, and protist components within the same sample [5]. Moreover, it is currently the most widely used sequencing technology and the one that generates the largest amount of publicly available data, which makes it especially valuable for the study of gut fungi.

Despite these technological advances, inconsistencies persist in the literature. For example, studies of the human gut mycobiome frequently report fungal taxa whose presence is unlikely to reflect true colonization of the gastrointestinal tract, as many of these organisms are environmental fungi or edible species derived from the diet and possess highly specialized nutritional requirements incompatible with gut growth [6], [7]. For instance, major studies such as the Human Microbiome Project have described as part of the “healthy” gut mycobiome a range of fungal taxa that are unlikely to represent true gut colonizers, including edible mushrooms such as *Agaricus bisporus* and *Ustilago maydis*, as well as plant-associated saprotrophs like *Alternaria* spp. and *Botrytis* spp. [2]. Likewise, an enterotype-based analysis of the human mycobiome reported the presence of additional edible genera, including *Auricularia* spp. and *Hypsizygus* spp., and even detected members of the phylum *Chytridiomycota* at approximately 3% relative abundance, despite these organisms being parasitic or saprophytic fungi with ecological and physiological requirements that are incompatible with sustained survival in the human gut [8].

This pattern also appears in disease-associated studies. Reports describe environmental fungi (*Mortierella* spp.) proposed as putative markers of metabolic disease, reduction of edible mushrooms (*Marasmius* spp., *Agaricus* spp.) in individuals with diabetes [9], and associations of dietary taxa such as *Agaricus* in pediatric-onset multiple sclerosis [10] or *Auricularia* in gestational diabetes and post–weight-loss interventions [11]. Also wood or plant-saprotrophic genera, such as *Tomentella* spp. and *Didymosphaeria* spp., whose occurrence in faecal samples is likely incidental and biologically irrelevant, have been associated to disease [12]. Collectively, these observations underscore the need to distinguish transient dietary or environmental passengers from gut colonizers when interpreting mycobiome data.

A second challenge relates to taxonomic classification. Widely used bioinformatic tools such as Kraken2 [13] and MetaPhlAn4 [14], were designed for bacteria and archaea, and while they can identify fungi, their accuracy for eukaryotic taxa is limited [15]. Dedicated tools such as MiCoP [16], FunOMIC [17], EukDetect [18], and HumanMycobiomeScan [19] have improved fungal detection, yet their performance depends heavily on the underlying database. A further obstacle is the lack of benchmarking and standardization: many fungal profiling methods are not consistently maintained, and gold standards for mycobiome analysis are still absent [15].

Relying solely on algorithm improvements is insufficient to reach a gold standard in the mycobiome analysis. Databases therefore play a critical role. Most pipelines do not allow users to customize the default database [14], [17], [18], which limits the possibility of constructing niche-specific catalogues. These have been shown to improve performance and reduce uncertainty in taxonomical identification [20], [21]. Moreover, available fungal databases often include incomplete, redundant, or contaminated assemblies, which can lead to spurious assignments and false-positive identifications [22], [23]. Recent work has highlighted the need for curated catalogues of fungal genomes specifically tailored to the human gut, ideally based on culture-dependent evidence to ensure biological relevance [24].

In this study, we developed FungiGutDB v1.0, a curated database composed exclusively of fungal species previously demonstrated to grow in the human gut, with the aim of reducing off-target taxonomic assignments in mycobiome characterization. We implemented this database within a publicly available pipeline to ensure reproducibility, accessibility, and interoperability in line with FAIR principles, and benchmarked its performance with simulated and culture-derived mock communities, as well as two independent human gut metagenomic cohort, in comparison with the standard reference databases.

This combined approach addresses current challenges in mycobiome research by providing both a high-quality database and a reproducible workflow optimized for the human gut ecosystem, reducing misclassifications due to the presence of environmental or contaminant fungi.

## Methods

### Workflow design and overview

The FungiGut workflow, built on Nextflow, a scalable workflow management system for reproducible and parallel bioinformatic analyses, has been designed following the FAIR principles (Findable, Accessible, Interoperable, and Reusable), with version-controlled code and explicit database snapshots. It comprises three main components (i) construction and indexing of a curated fungal database (FungiGutDB v1.0); (ii) read preprocessing, quality filtering and removal of host and bacterial DNA; and (iii) fungal profiling by MiCoP alignment and filtering to derive species-level results (Figure 2). Performance was evaluated on simulated mock communities and on two real-world human gut sample cohorts.

**Figure 1:**
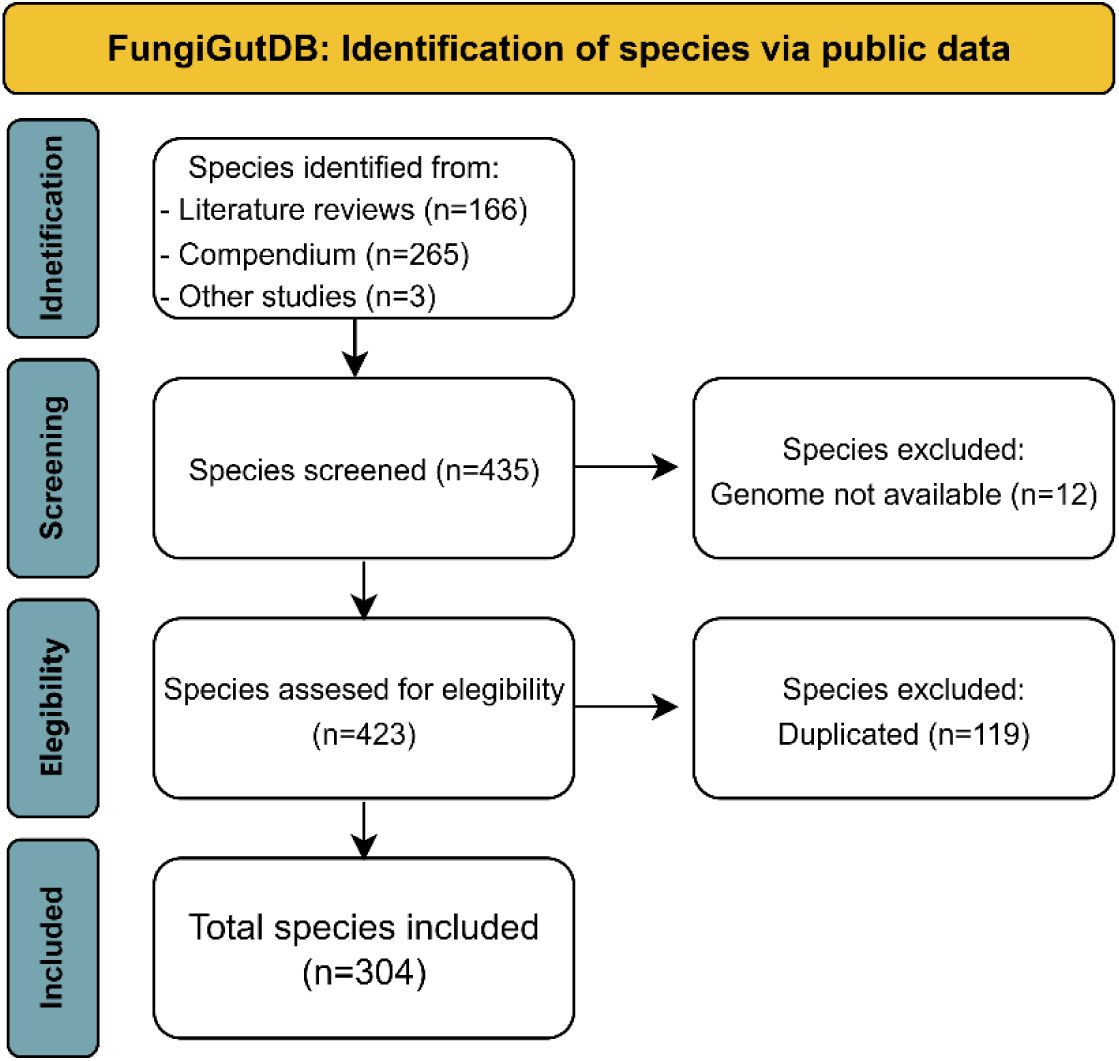
Species identification workflow for FungiGutDB v1.0. Overview of the steps used to select and validate fungal species included in the database.

**Figure 2:**
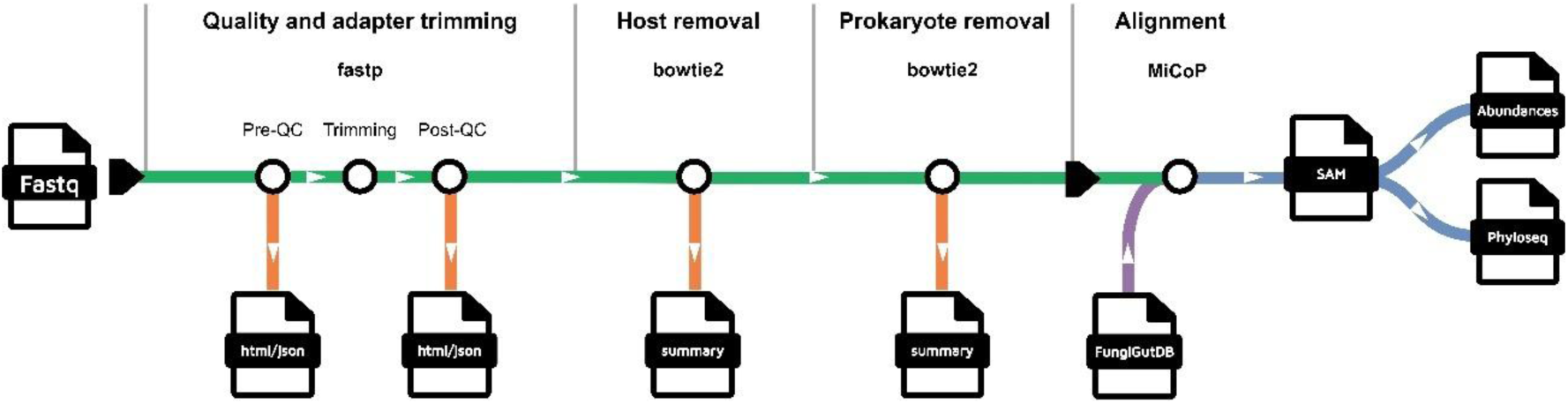
FungiGut nextflow workflow.

#### (i) Fungal database

##### Sources and species list

The studies used for database species selection were retrieved from public sources. The core species database relies on a recent study that compiled a genomic compendium of fungal species from the human gut and also provided an NCBI list of human-associated fungi, both of which were incorporated into FungiGutDB [24]. Specifically, the genomic compendium corresponds to the cultivated gut fungi (CGF) catalogue, which consists of 760 fungal genomes isolated from the faeces of healthy individuals. This catalogue includes 206 species across 48 families, among them 69 species previously uncharacterized, thereby substantially expanding the known diversity of gut-associated fungi. In addition, three reviews which provide culture-based studies of human gut fungi were identified and included. The most recent of these reviews covered the literature up to 2017 [3], [7], [25]. To extend coverage, different studies published between 2017 and 2025 that employed culture-dependent methods for fungal identification were examined; however, only three of these contributed additional species to the database [26], [27], [28] (Supplementary Table S1, Panel 2). Species lacking available assemblies in GenBank or RefSeq were excluded (Figure 1).

Fungal genome assemblies were retrieved from NCBI Datasets (downloaded 2025-08-01). The database is intended to be dynamic and can be versioned and updated as new assemblies become available. The snapshot used in this study (FungiGutDB v1.0) can be used as the default reference within the pipeline, while an update option allows rebuilding the database from the curated species list to incorporate newly released assemblies.

##### Assembly selection

To minimize assembly bias, a hierarchical selection scheme per species was applied: Refseq (if available) > reference genomes (if available) > complete genomes > chromosome > scaffold/contig assemblies with ties broken by the lowest number of scaffolds, to obtain the best assembly metrics and qualities. A single assembly per species was included, to avoid redundancy. To filter quality of assemblies and very short fragments, scaffolds shorter than 500 bp were removed using seqkit (v2.10.0) [29].

##### Taxonomic metadata and indexing

Accession-to-taxon metadata were compiled from NCBI Taxonomy (downloaded on 2025-08-01 for FungiGutDB v1.0), following the structure needed for MiCoP method (accession ID, accession length, taxid, full taxonomy lineage).

The final merged fasta after selection and filtering was indexed with *BWA-index* (0.7.19-r1273) [30] for downstream alignment. This indexed reference (FungiGutDB v1.0) is distributed alongside the workflow as the default database.

#### (ii) Read preprocessing and decontamination

##### Quality control and trimming

Reads are processed with *fastp* (v0.25.0) [31]: adapter auto-detection, per-base quality trimming (Q≥20), removal of reads with length < 50 bp, and filtering of reads containing >5% ambiguous bases. Pre- and post-QC reports are generated to verify read quality.

##### Host and bacterial read removal

To suppress false fungal matches from residual host or bacterial DNA, reads are sequentially mapped to (i) the *human reference* (GRCh38) and (ii) the *Unified Human Gastrointestinal Genome* (UHGG) catalogue (v2.0.2) [21], using *Bowtie2* (v2.5.4) [32], retaining only unmapped reads for fungal profiling. Mapping statistics (input reads, % host-removed, % bacteria-removed, remaining reads) per sample are recorded.

#### (iii) Fungal profiling: alignment and abundance estimation

Fungal profiling follows a MiCoP alignment approach. Briefly, bacteria-filtered reads are aligned to FungiGutDB with bwa-mem (0.7.19-r1273). Then, alignments are filtered using four parameters: minimum mapped length (*min_map*), maximum edit distance (*max_ed*), minimum identity (*pct_id*), and minimum reads to report a taxon (*read_cutoff*). Unless stated otherwise, defaults are *min_map*=100, *max_ed*=1, *pct_id*=0.99, *read_cutoff*=10. Alignments lacking a valid accession mapping in accession2info.txt are discarded.

As in MiCoP, multi-mapped reads are handled by probabilistic reassignment: in default mode, they are redistributed and then normalized by genome length before conversion to relative abundance, whereas in raw count mode they are assigned as whole reads to taxon following the same proportional rule without length normalization.

Final outputs include species-level abundances/read count tables per sample and an RDS object compatible with *phyloseq R package* [33] for downstream analyses.

### Datasets

#### *In silico* mock communities

To assess detection accuracy under different noise regimes, 40 simulated metagenome samples were computationally generated with CAMI-based software *MeSS* (v0.11.0) [34] using Illumina-like error profiles (2×150 bp) and default settings specified. Species were sampled at random from the FungiGutDB (*Supplementary Table S2)*.

Two scenarios were simulated (20 samples per set):

1. Mixed bacterial/fungal (Set1): 15 fungal species + 5 bacterial species; 95% of reads bacterial and 5**%** fungal. Each sample has ~5G bases count. This scenario was run with and without bacterial filtering to quantify the benefit of the (UHGG) step.
2. High fungal complexity (Set2): 40 fungal species; total bases count per sample was ~1G.

#### Public mock communities

Two mock communities from previous studies were used for validation. Unlike the *in silico* mock communities, these underwent an actual sequencing process, making them closer to real metagenomic data.

1. The first culture-derived mock community comprised two fungal species [35]. Specifically, the commercially available community (Catalog No. D6300, Zymo Research) contains cells from eight bacterial species (each representing 12% of the community) and two yeast strains (each contributing 2%, representing a 1:1 proportion).
2. The second is a DNA-derived mock community, generating paired-end Illumina shotgun metagenomic sequencing from pooled DNA. This mock is composed of 44 fungal strains representing 39 different species [36].

#### Real-world datasets

Two publicly available human gut metagenomic cohorts were analysed to assess FungiGut performance on real-world data including healthy and a clinically non-healthy population, allowing evaluation in fungal detection in a clinically distinct, gut environments.

##### Healthy population

The first cohort consisted of 60 stool metagenomes from healthy controls recruited within the MetaCardis project [37]. In this study, healthy controls are defined as metabolically healthy adults without diagnosed cardiometabolic, gastrointestinal or inflammatory diseases, and not undergoing any pharmacological treatment known to alter gut microbiome composition. These individuals serve as a baseline reference population widely used in microbiome analyses within the MetaCardis consortium.

##### Non-healthy population

The second cohort included 39 stool metagenomes from patients with non-responsive celiac disease (NRCD) [38]. NRCD patients are clinically characterised by persistent gastrointestinal symptoms despite long term adherence to a strict gluten-free diet and evidence of altered gut microbiome structure, highlighted in the original study.

### Performance evaluation and statistical analysis

#### Evaluation metrics

For synthetic *in silico* and culture-derived mock communities with known composition, species level identification performance was evaluated in terms of F1-score, computed using true positives (expected and detected), false positives (detected but not expected), and false negatives (expected but not detected). Relative abundance differences between the expected and observed were measured using root mean square error (RMSE).

#### Statistical analyses

Downstream analyses were performed in *R* (v4.4.3), handling compositional fungal data with *phyloseq* (v1.50.0). The phyloseq object was filtered differently depending on the dataset. For *in silico* mock communities, only species with relative abundances greater than 0.1% were retained. For public mock communities and real-world datasets, a more permissive threshold was applied, accounting for the higher variability and noise typical of experimental metagenomic data. Species with relative abundances below 0.01% were excluded.

Absolute abundances were transformed to compositional when required, using *microbiome R package* (v1.5.0) [39]. Alpha-diversity was assessed using Chao1, Shannon and Simpson indices and beta-diversity was performed with weighted UniFrac distance.

Where applicable, non-parametric Wilcoxon tests were used for group comparisons, and p-values were adjusted for multiple testing using the Benjamini-Hochberg method.

#### Batch-effect correction

To account for technical variability between real-world datasets, batch correction was performed using the *MMUPHin R package* (v1.20.0) [40]. Adjustment was applied with the study of origin specified as the batch variable. The impact of correction was assessed by quantifying the variance in fungal community structure explained by study before and after adjustment. This evaluation was conducted using permutational multivariate analysis of variance (PERMANOVA, 999 permutations) implemented with de *adonis2* function from the *vegan R package* (v2.6-10) [41]. A reduction in variance attributable to study after correction was interpreted as effective mitigation of batch effect related effects.

## Results

### FungiGutDB a comprehensive curated database for the human gut mycobiome

The initial release of FungiGutDB (v1.0) comprises 304 taxa previously informed in culture-dependent human studies (Supplementary Table S1). Species are distributed across five phyla reported in culture-based surveys: *Ascomycota* (~79.27%), *Basidiomycota* (~12.82%), Mucoromycota (~6.25%), Microsporidia (~1.31%), and *Zoopagomycota (<1%)* (Figure 3). The database includes 124 genera; the most abundant genera in terms of number of species are *Aspergillus* (35 species, ~11%) followed by *Penicillium* (10 species, ~8%), and *Candida* (17 species, ~5%).

**Figure 3.**
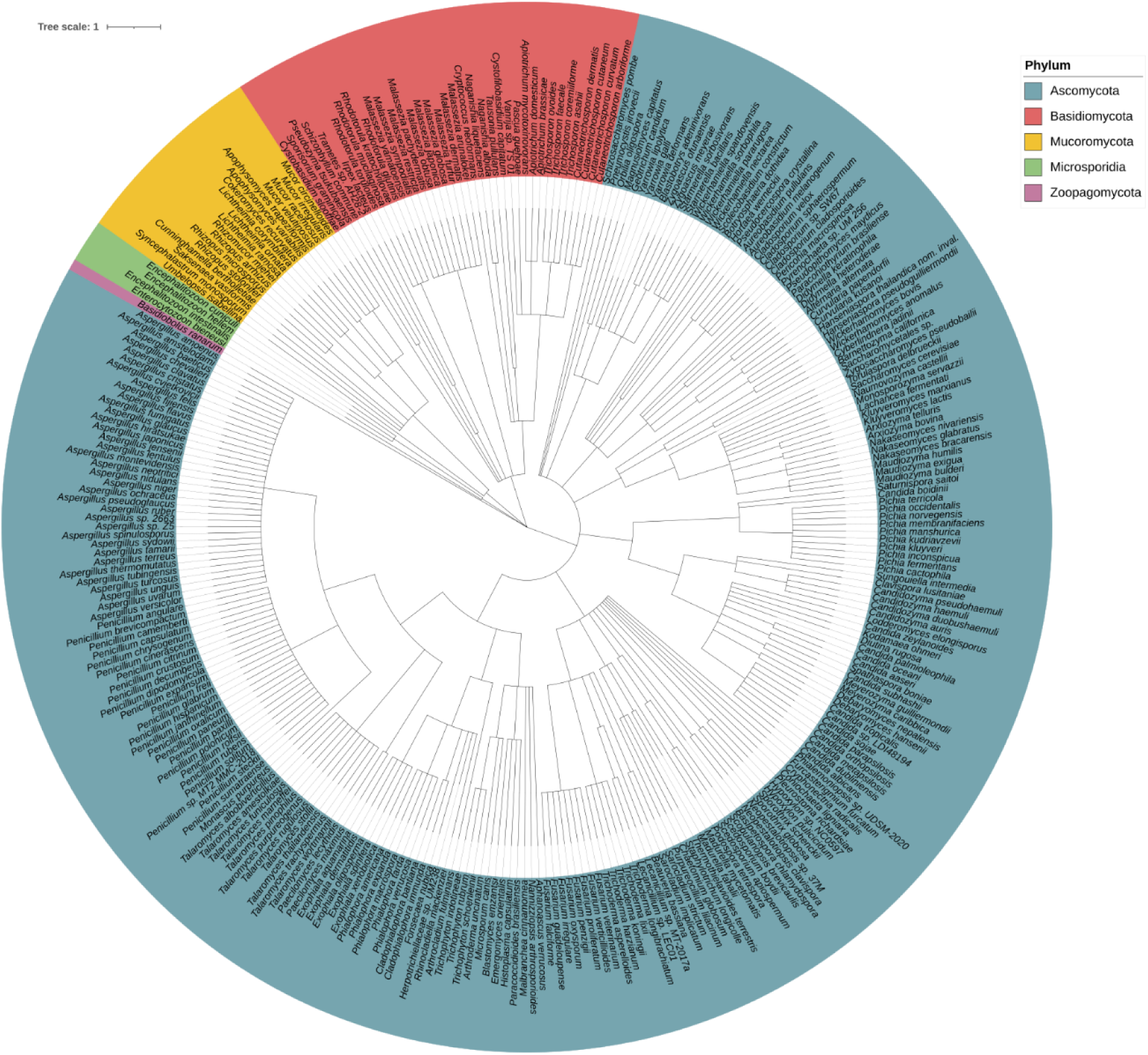
Species distribution in FungiGutDB v1.0. The database includes 304 distinct species from five phyla: *Ascomycota* (~79.27%), *Basidiomycota* (~12.82%), Mucoromycota (~6.25%), Microsporidia (~1.31%), and *Zoopagomycota (<1%)*.

Assemblies were primarily sourced from RefSeq (~37%) with the remainder from GenBank (~63%). In both subsets, most entries were tagged as reference assemblies. Assembly accessions, taxonomy IDs, assembly level, and scaffold count metrics are listed in Supplementary Table S1.

### FungiGut shows high performance on synthetic mock communities

We implemented MiCoP for taxonomic assignment in the FungiGut pipeline, as it has been reported to outperform other tools (Kraken2, MetaPhlAn4) and, unlike EukDetect and FunOMIC supports the integration of custom databases [10]. Database used for this purpose was FungiGutDB v1.0.

Set 1 was designed to emulate metagenomic samples where most of the genetic content originates from bacteria, allowing evaluation of the impact of bacterial background on fungal detection. To assess this effect, samples were profiled both before and after filtering against the UHGG database. FungiGut achieved improved performance after prokaryotic filtering, in terms of false positives and F1-score (0.727 in Set1-nofilter vs. 0.827 in Set1-filtered; Wilcoxon, BH-adjusted p = 2.31e07), while no significant changes were found in RMSE (Wilcoxon, BH-adjusted p = 9.09e-2) (Figure 4).

**Figure 4:**
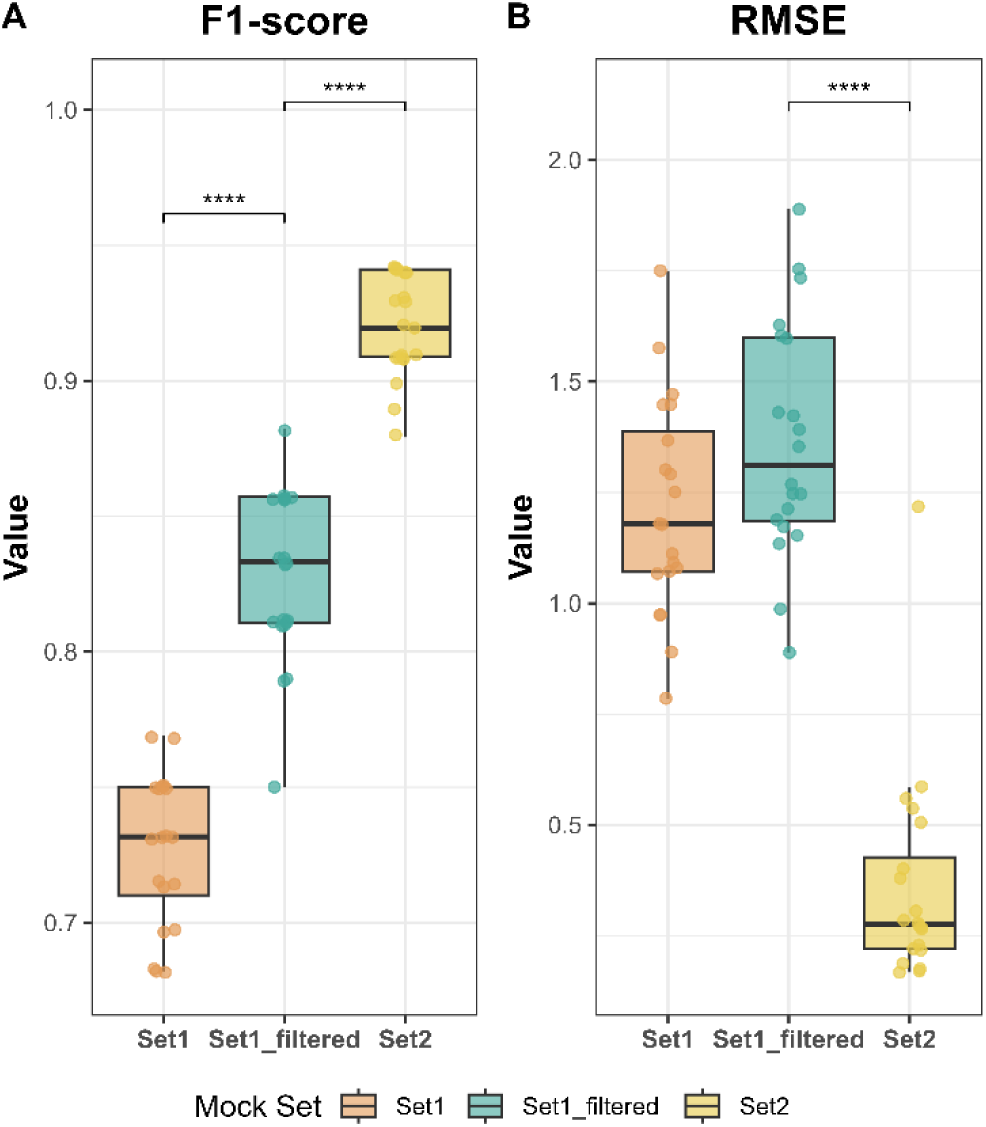
Synthetic mock analysis. A) F1-score and, B) RMSE for Set1, Set1_filtered (after removing prokaryotic reads), and Set2, which contains a higher fungal abundance and does not require prokaryotic filtering.

Set 2 consisted of synthetic fungal communities with increased species diversity and no bacterial background, enabling evaluation of performance under conditions of higher fungal diversity but absence of prokaryotic interference. Under these conditions, the pipeline reached an overall F1-score of 0.919 and RMSE of 0.3617.

Comparing the filtered Set 1 and Set 2 provides a complementary evaluation of two key determinants of profiling accuracy: background bacterial interference and fungal community complexity. Set 2 displayed significantly higher F1-score (Wilcoxon, BH-adjusted p = 1.26e−07) and significantly lower RMSE values (BH-adjusted p = 1.91e−07), indicating that once bacterial background is minimized, increased fungal diversity does not compromise taxonomic accuracy (Figure 4).

These results demonstrated robust taxonomic performance across all mock scenarios, with prokaryotic filtering increasing F1-score by ~14%.

### FungiGut pipeline outperforms standard MiCoP database

The pipeline performance was tested against MiCoP with the standard database (RefSeq) using two publicly available mock communities. All the FungiGut pipeline steps were applied to the samples. Host and UHGG sequences decontamination was performed after a quality trimming. For comparison, the MiCoP tool was run with the standard database using the quality trimmed reads. This evaluation aimed to assess the impact of using the pipeline and FungiGutDB v1.0 on culture derived mock communities.

For the mock community (Catalog No. D6300), FungiGutDB v1.0 recovered the two expected species in the sample. *Cryptococcus neoformans* represented 70.05% of the community, and *Saccharomyces cerevisiae* 29.34%. Three additional taxa were detected at very low relative abundances: *Cladosporium velox* (0.28%), *Candida subhashii* (0.20%), and *Geotrichum candidum* (0.13%).

Using the standard MiCoP database, the two target species, *C. neoformans* and *S. cerevisiae*, were also recovered, with relative abundances of 66.71% and 28.82% respectively. However, additional species were reported with higher relative abundances compared to FungiGut database. Among them were *Wickerhamomyces ciferrii* (1.49%), *Enterocytozoon bieneusi* (1.45%), *Cryptococcus gattii* (0.85%), *Diplodia corticola* (0.26%), *Kwoniella pini* (0.25%), and *Candida auris* (0.16%).

The second mock community contained DNA from 44 fungal strains representing 39 distinct species. In this analysis, we compared the two databases by assessing the presence or absence of taxa and recording the false positives detected. Using the FungiGutDB v1.0, 30 species were identified, including 27 true positives and 3 false positives. In contrast, the MiCoP standard database recovered 19 species, of which 17 were true positives and 2 were false positives. Detailed results, including the full list of species detected and expected in each analysis are available in Supplementary Table S3.

The results show that both approaches correctly identified the target species in the mock datasets whereas neither method recovered the expected proportion of species 1:1 in the first mock, while FungiGut pipeline, and already mentioned database version, recovered higher number of expected taxa and fewer spurious detections across the tested mock communities.

### FungiGut performance on real world data

The FungiGut workflow and FungiGutDB v1.0 were applied to two independent cohorts: healthy controls from the MetaCardis project (*n* = 60, single-end reads) and a set of patients with non-responsive celiac disease (NRCD, *n* = 39, paired-end reads). For comparison, the same datasets were analysed with MiCoP using its standard fungal reference database (RefSeq). In both cases, the resulting phyloseq objects were filtered to retain taxa present at >0.01% abundance in at least one sample. Detailed species-level results for both cohorts are provided in Supplementary Table S3.

#### MetaCardis cohort

In the MetaCardis healthy controls (*n* = 60), FungiGut detected a total of 54 species distributed across 34 genera. By comparison, analysis with the MiCoP standard database recovered only 16 species belonging to 13 genera.

The most abundant species in the FungiGut profiles was *S. cerevisiae* (mean relative abundance = 69.0%), followed by *G. candidum* (12.45%), *Penicillium camemberti* (8.0%), *Debaryomyces hansenii* (3.2%), and *Pichia kudriavzevii* (1.4%). Additional species consistently present included *Candida albicans* (1.0%), *Kluyveromyces lactis* (0.9%), *Pichia occidentalis* (0.6%), *Torulaspora delbrueckii* (0.4%), and *Candida parapsilosis* (0.3%).

In contrast, MiCoP with the standard database reported markedly fewer taxa. The most abundant species was again *E. bieneusi* (78.4%), followed by *S. cerevisiae* (14.1%) and *W. ciferrii* (3.8%). Other taxa included *D. hansenii* (0.8%), *Sordaria macrospora* (0.8%), *P. kudriavzevii* (0.6%), *K. lactis* (0.4%), *Saccharomyces eubayanus* (0.2%), *Grosmannia clavigera* (0.1%), and *Kluyveromyces marxianus* (0.07%).

Frequency patterns also highlighted differences between the two databases. With FungiGut, *S. cerevisiae* was detected in nearly all samples (58 of 60), together with *G. candidum* (41 samples) and *P. camemberti* (24). Other recurrent taxa included *D. hansenii* (14 samples), *K. lactis* and *T. delbrueckii* (8 each). In contrast, MiCoP profiles were again dominated by *E. bieneusi* (60 of 60 runs), followed by *S. cerevisiae* (56 samples), *W. ciferrii* (30), and *S. macrospora* (19).

#### NRCD cohort

Using FungiGut, the NRCD samples yielded a total of 72 distinct species belonging to 43 genera. In contrast, analysis with the MiCoP standard database identified 48 species distributed across 33 genera. The higher number of taxa recovered with FungiGut was accompanied by a broader representation of known gut-associated fungi.

The most abundant species detected with FungiGut included *S. cerevisiae* (mean relative abundance = 52.9%), followed by *P. camemberti (*4.4%), *G. candidum* (4.3%), *Fusarium proliferatum* (3.9%), *Barnettozyma californica* (3.8%). Other taxa consistently detected were *Malassezia restricta* (3.4%), *D. hansenii* (3.1%), *Aspergillus niger* (2.8%), *Malassezia pachydermatis* (2.8%), and *Sungouiella intermedia* (2.5%).

By contrast, the MiCoP standard database reported a markedly different taxonomic profile. The most abundant taxon was *E. bieneusi* (81.9%), followed by *S. cerevisiae* (7.3%) and *W. ciferrii* (5.1%). Additional taxa included *Debaryomyces hansenii* (1.5%), *S. macrospora* (1.4%), *G. clavigera* (0.3%), and lower-abundance species such as *M. pachydermatis*, *A. niger*, *Aspergillus flavus*, and *Malassezia globosa*.

When considering prevalence across samples, *S. cerevisiae* was present in 30 out of 39 NRCD samples according to FungiGut, followed by *G. candidum* (17 samples), *F. proliferatum* (12), and several taxa such as *A. niger*, *D. hansenii*, and *P. camemberti* each appearing in 7 samples. In contrast, MiCoP profiles were dominated by *E. bieneusi*, detected in all 39 runs of the dataset, together with *W. ciferrii* (27 samples) and *S. macrospora* (23), while *S. cerevisiae* was found in only 17 samples.

#### Cohort comparison

To assess the resolution of both approaches in detecting community level variation, NRCD and MetaCardis cohorts were compared using beta and alpha diversity metrics.

With the MiCoP standard database, no significant differences between groups were observed. Beta-diversity analysis based on weighted UniFrac distances yielded an *R^2^* of 0.0139 (F = 1.39, p = 0.228), indicating no clear separation between NRCD and non-celiac samples (Figure 5C). Likewise, alpha-diversity indices (Chao1, Shannon, Simpson) showed no significant differences between groups (all p > 0.05; Figure 5D).

**Figure 5:**
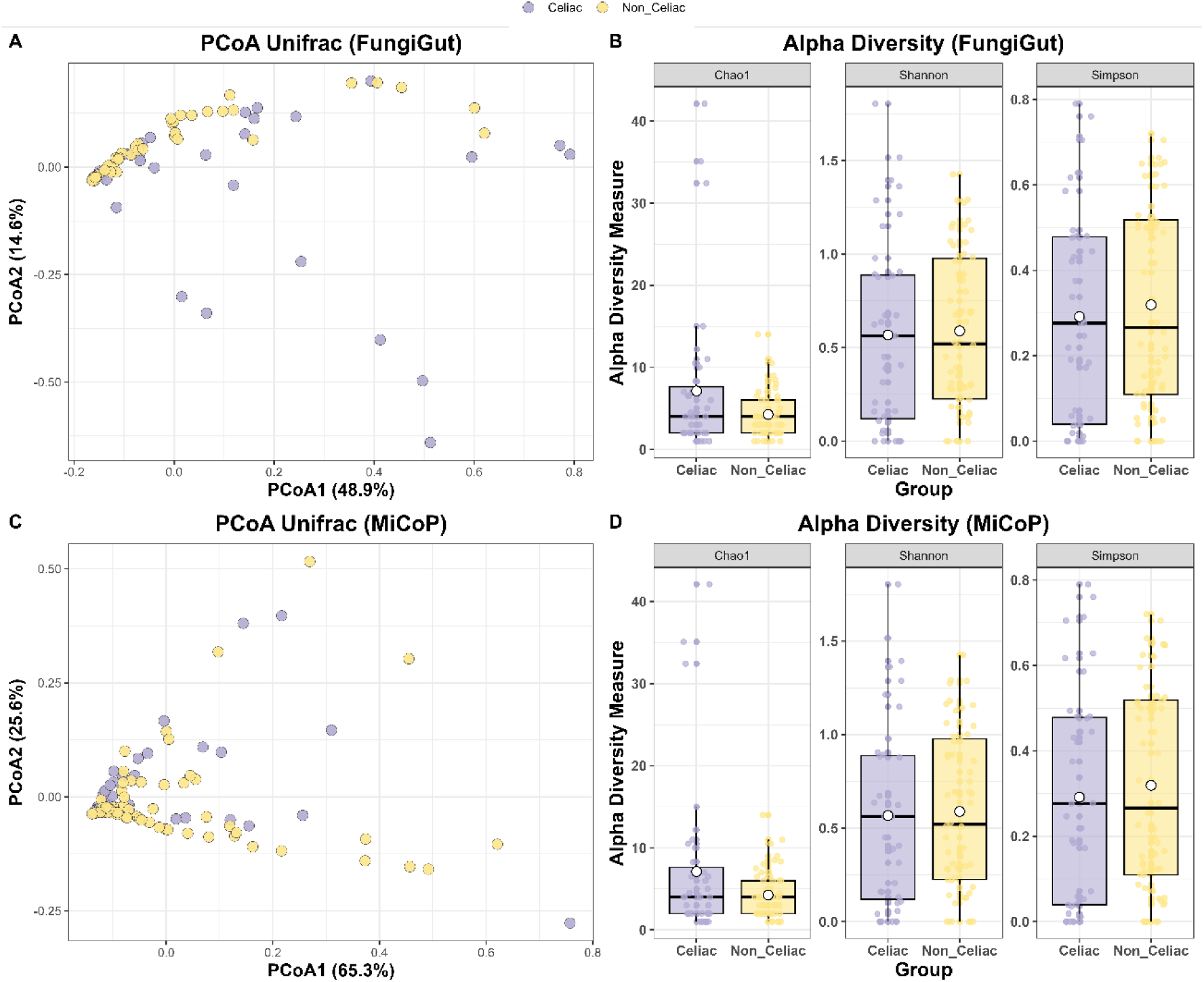
Gut mycobiome community analysis. Alpha diversity (A, C) and UniFrac PCoA (B, D) comparing Celiac and Non-Celiac individuals. Results are shown for the FungiGut workflow (A–B) and the MiCoP approach (C–D).

By contrast, FungiGut revealed clearer community-level differences. Weighted UniFrac analysis showed a significant separation between NRCD and controls (*R^2^* = 0.0428, F = 4.25, p = 0.004) (Figure 5A). Although alpha-diversity indices (Chao1, Shannon, Simpson) did not differ significantly (all p > 0.05; Figure 5B), the beta-diversity signal suggests that the curated fungal database provides a higher resolution of inter-group variation.

## Discussion

The gut mycobiome plays important functions in the maintenance of homeostasis and its importance in the development of various diseases has been recognized; however, its correct characterization remains somewhat neglected [15], [42]. Improved tools for accurate taxonomic identification are urgently needed, as previous studies have reported fungal species that are nutritionally incapable of growing in the human gut, such as plant pathogens or environmental and edible fungi, as part of the normal mycobiome [8], [43].

A recent study made an important contribution by generating a curated, fungi-specific database in which human-derived contaminants and other sources of noise were removed, drawing on sequences from major fungal repositories (Ensembl Fungi, FungiDB, MycoCosm, and NCBI RefSeq) [44]. While this effort is significant, the resulting resource is not restricted to human-associated fungi. From our perspective, this may perpetuate a critical source of bias: the inclusion of environmental, edible, and foodborne fungi that cannot grow in the human gastrointestinal tract, yet are disproportionately represented in public fungal sequencing databases. Such taxa can confound taxonomic profiles and inflate biologically implausible assignments.

To address these limitations, we curated a database specifically focused on the intestinal mycobiome, including only those fungi that have been previously described in the literature or that are biologically plausible colonizers or transient members of the human gut in physiologically reasonable abundances. Our database was integrated into a state-of-the-art pipeline for fungal profiling [15], [16], thereby leveraging existing resources developed by other research groups. Additionally, we implemented this workflow in Nextflow, ensuring reproducibility and accessibility through an open and flexible framework.

Our results demonstrate that FungiGut accurately characterizes the intestinal mycobiome from shotgun metagenomic data using standard preprocessing steps. When compared with the standard MiCoP pipeline, FungiGut achieved a lower false-positive rate in mock communities, detecting only three low-abundance contaminants (0.61% abundance), whereas MiCoP identified six false positives species (4.46% abundance), including the identification of an emerging clinical pathogen: *Candida auris*, currently listed among the World Health Organization’s priority fungal pathogens [45], [46]. These results not only introduce background noise by detecting a species that is unlikely to be truly present but could also trigger unnecessary alarm if such a pathogen was reported in a clinical or hospital environment.

When applied to real datasets, FungiGut not only recovered the intestinal mycobiome composition but also yielded biologically meaningful results. In healthy subjects, FungiGut identified *S. cerevisiae* as the most abundant fungal species, followed by *G. candidum* and several diet-associated fungi present in dairy fermented products and cheese, such as *P. camemberti, D. hansenii, K. lactis,* and *P. kudriavzevii* [47], [48], [49], [50]. In contrast, MiCoP reported lower species diversity, characterized by a reduced proportion of *S. cerevisiae* and a marked dominance of *E. bieneusi,* a microsporidian parasite of humans and animals transmitted through contaminated food or water [51]. MiCoP also detected environmental taxa *W. ciferrii* (found in pods and plant exudates [52]), *S. macrospora* (found in soil and dung [53]), and *G. clavigera* (a tree pathogen associated with bark beetles [54]), and fewer diet-associated species [55] (*K. lactis, P. kudriavzevii, D. hansenii, S. eubayanus, K. marxianus*).

In patients with non-responsive celiac disease (NRCD), FungiGut identified *S. cerevisiae* as the most abundant fungus. Several opportunistic pathogens were also detected, including yeast associated with fungemia in immunocompromised patients [56], [57], [58] (*M. restricta, M. pachydermatis, R. mucilaginosa*) and molds causing invasive fungaemia [59], [60] (*F. proliferatum, A. niger*). Dietary-associated fungi from dairy and fermented products (*D. hansenii, P. camembert*i) and olive oil (*B. californica* [61]) were also present. Conversely, MiCoP detected *E. bieneusi* at over 70% relative abundance, an unexpectedly high prevalence, given that this parasite is typically associated with infections in immunocompromised patients [62]. MicoP further identified several environmental fungi such as *W. ciferrii, S. macrospora, G. clavigera and Aspergillus flavus* [63], and a lower detection of opportunistic pathogens previously involved in invasive fungaemia (*M*. *pachydermatis, A. niger*) and dermatitis [64] (*M*. *globos*a) including also the multi-drug-resistant pathogen *C. auris*.

Our curated database thus restricted identification to fungi previously isolated from the human gut, whether as commensals, opportunistic pathogens, or dietary transients, while substantially reducing or eliminating environmental or plant-associated contaminants. FungiGut identified *S. cerevisiae* and other commensal or dietary fungi in healthy individuals, whereas in NRCD patients, a condition characterized by persistent intestinal symptoms and inflammation, it revealed a higher prevalence of opportunistic pathogens potentially contributing to disease pathophysiology alongside bacterial dysbiosis. A non-gut specific database, by contrast, fails to capture these biologically relevant variations in gut mycobiome composition, instead introducing environmental noise and misleading identifications.

Our study has several limitations. First, we did not implement a DNA extraction method specifically optimized for fungi. Fungal cell walls, composed mainly of chitin, glucans, mannans, and glycoproteins, are difficult to lyse and can substantially affect DNA yield and quality [3]. Using standard protocols may therefore limit the recovery of fungal sequences. Because our workflow begins with previously generated sequencing data, we cannot account for extraction-related biases.

Additionally, our database includes some dietary and allochthonous fungi (*D. hansenii, P. camemberti*, *Saccharomyces* spp.). Determining whether these taxa exert a meaningful biological role in the gut remains challenging. Despite their limited ability to grow at 37°C or to establish true intestinal colonization, these species have been repeatedly isolated in studies and, as frequent dietary components, may contribute metabolites or postbiotic compounds [3], [6]. However, their precise functional relevance cannot be determined from our current analyses. Similarly, the presence of certain opportunistic pathogens, particularly in the NRCD cohort, may reflect a dysbiotic state, yet their true pathogenic potential and physiological impact cannot be resolved using these methods.

Further population-based studies are needed to clarify the role of these fungi across diverse host contexts, including immunocompromised individuals, patients with hematopoietic disorders, and neonatal populations [3], [65]. Functional investigations are also necessary to elucidate interkingdom interactions and the potential metabolic or immunological roles of these taxa. To date, our work cannot conclusively determine whether the detection of fungal species, especially diet-associated taxa, reflects active and functional participation in the intestinal mycobiome or merely transient colonization without physiological relevance. Nevertheless, their detection is biologically meaningful, as these species are neither environmental contaminants nor organisms incapable of survival within the human gastrointestinal environment.

In light of the current challenges in the field, namely experimental noise, taxonomic misidentifications, and biologically implausible interpretations that collectively undermine the credibility of intestinal mycobiome research, FungiGut emerges as a framework aimed at restoring biological consistency and interpretative accuracy to mycobiome studies, firmly grounded in both clinical and ecological mycology.

## Acknowledgements

Figure 3 was generated using iTOL: Interactive Tree Of Life (<u>Letunic I and Bork P</u> (2024) Nucleic Acids Res doi: 10.1093/nar/gkae268 *Interactive Tree of Life (iTOL) v6: recent updates to the phylogenetic tree display and annotation tool)*.

## Availability of data and material

Sequence files for all human samples used in this study are available in ENA under the accession numbers PRJEB65879 and PRJEB41311, PRJEB38742 and PRJEB37249. Culture-derived public mock communities have been downloaded from PRJNA1000750 and SRX10705695. A full record of all data analysis and original R scripts are available in https://github.com/diegocoleto7/FungiGut-Paper. Pipeline scripts are available in https://github.com/diegocoleto7/FungiGut. The FungiGutDB v1.0 database snapshot is archived at Zenodo: https://doi.org/10.5281/zenodo.17581472.

## Authors’ contributions

D. C-C, conceptualization, data curation, formal analysis, software, writing - original draft preparation; B. L-P, visualization, writing - review & editing; A P-C, validation, writing - review & editing; NC-R, validation, writing - review & editing; L. C-G, validation, writing - review & editing; A. M-S: resources, writing - review & editing; E CSP, writing - review & editing, funding acquisition, supervision, L. J. M-Z: conceptualization, methodology, funding acquisition, writing - original draft preparation

